# Network-based amyloid-β pathology predicts subsequent cognitive decline in cognitively normal older adults

**DOI:** 10.1101/2024.12.10.627818

**Authors:** Hengda He, Qolamreza R Razlighi, Yunglin Gazes, Christian Habeck, Yaakov Stern, Alzheimer’s Disease Neuroimaging Initiative

## Abstract

The deposition of amyloid-β (Aβ) protein in the human brain is a hallmark of Alzheimer’s disease and is related to cognitive decline. However, the relationship between early Aβ deposition and future cognitive impairment remains poorly understood, particularly concerning its spatial distribution and network-level effects. Here, we employed a cross-validated machine learning approach and investigated whether integrating subject-specific brain connectome information with Aβ burden measures improves predictive validity for subsequent cognitive decline. Baseline regional Aβ pathology measures from positron emission tomography (PET) imaging predicted prospective cognitive decline. Incorporating structural connectome, but not functional connectome, information into the Aβ measures improved predictive performance. We further identified a neuropathological signature pattern linked to future cognitive decline, which was validated in an independent cohort. These findings advance our understanding of how Aβ pathology relates to brain networks and highlight the potential of network-based metrics for Aβ-PET imaging to identify individuals at higher risk of cognitive decline.

## Introduction

Amyloid-β (Aβ) pathology is recognized as one of the earliest initiating events in Alzheimer’s disease (AD) progression. As proposed by the Amyloid-β Cascade Hypothesis (ACH)^1,2^, abnormal accumulation of Aβ in the brain has been linked to a cascade of pathological events, including tau tangle formation and spread^3,4^, inflammation^5^, gray matter atrophy^6^, neuronal dysfunction and network disruption^7,8^, and ultimately cognitive and functional impairment^9^. Aβ deposition is commonly observed in the brain in older adults who do not show significant cognitive impairment^10^. However, individuals with elevated Aβ pathology are at an increased risk of developing dementia in the future^11^, and are considered to be in the preclinical phase of AD^12^. In this study. we developed network-based amyloid-β pathology (NAP) measures that incorporate subject-specific connectome information. We then examined how these neuropathology measures, assessed at baseline, are associated with cognitive changes over the subsequent years.

The literature presents heterogeneous findings regarding the relationship between Aβ and cognition. The most consistent associations have been observed in episodic memory^13–17^. However, the reported relationships between Aβ and cognition are often weak^15,18^, and the results for non-memory cognitive domains are inconsistent. Some studies that assessed global cognition, combining measures across multiple cognitive domains, yielded slightly larger effect sizes, but such effects remain moderate^15^.

In the clinical settings, Aβ deposition is quantified as a dichotomous classification and is measured globally, covering the entire cerebral cortex. However, the organization of brain network systems has been proposed as an important mediating factor in the relationship between neuropathology and clinical symptoms^19–21^. Therefore, effective metrics of early Aβ pathology that consider its spatial spread are needed, and may potentially facilitate earlier recognition and intervention^22^.

Aβ deposition is spatially heterogeneous, especially in the preclinical phase of AD. Thus, some studies suggested that regional and continuous measures of Aβ deposition provide additional value in the early identification of participants at risk of transitioning from preclinical to clinical AD^14,24,25^. Early deposition has been shown to concentrate in the default mode network (DMN) and other highly connected regions^7,23^. By exploring the spatial pattern and regional-temporal evolution of Aβ deposition, studies have suggested diverse pathological staging patterns^26–28^. Moreover, recent evidence indicates that early Aβ deposition patterns vary across individuals^25,29^. These spatially heterogenous patterns may reflect the accumulation of neuropathology in distinct sets of brain networks, depending on the individualized connectome, and different networks may potentially be associated with distinct clinical symptoms. These findings highlight the need for integrating connectome and network information in Aβ-cognition studies.

In this study, we characterized the mediating role of the brain network connectome on the Aβ- cognition relationship by employing network-based neuropathological measures of Aβ. We reasoned that these measures would provide a more comprehensive metric to capture the complex relationship between neuropathology and cognitive function. We developed NAP measures by incorporating connectome information that consider both initial regional Aβ burden and its deposition within individualized connectomes. We then examined how these patterns, measured at baseline, are associated with cognitive changes over subsequent years. To improve the generalizability of our findings, we employed a cross-validated machine learning predictive model. We evaluated the performance of the NAP measure in predicting subsequent longitudinal cognitive decline, comparing it to the predictive performance of the regional amyloid-β pathology (RAP) measure, where RAP was measured as standardized uptake value ratio (SUVR) in the positron emission tomography (PET) imaging of Aβ pathology. We hypothesized that integrating connectome information would improve the prediction of cognitive decline. We derived a neuropathological signature of cognitive decline which provides insights into the regional-specific and network-level contributions of Aβ neuropathology to cognitive deterioration. We then sought external validation in the second cohort to confirm the neuropathological signature’s association with future cognitive decline, and demonstrate its generalizability. These results advance our understanding of Aβ pathology within brain network systems and offer a framework for future studies on Aβ deposition and cognition that incorporate personalized connectome profiles and brain networks analyses. The proposed NAP measures could potentially better reflect individual risk of future cognitive decline.

## Results

Participants were recruited through our ongoing cognitive reserve and reference ability neural network (CogRes/RANN) longitudinal study^30^. All participants were cognitively normal at both baseline and follow-up; they were screened for dementia and mild cognitive impairment (MCI) using the Dementia Rating Scale (DRS)^31^. A minimum score of 135 was required for the DRS assessment. Here, we included eighty-five cognitively normal older adults (Table 1; More details on participants inclusion criteria and demographic information are in the Methods section). The interval between baseline and follow-up neuropsychological assessments ranges 3 to 7 (4.51 ± 0.68, mean ± SD; SD, standard deviation) years. A subset of the neuropsychological assessments was used to examine participants’ cognition in four domains: episodic memory, vocabulary, processing speed, and fluid reasoning. Cognitive composite scores for each cognitive domain were generated and normalized by subtracting the baseline sample mean and dividing by the baseline sample standard deviation. An average of the four z-scores of all four cognitive domains was calculated to yield a global cognitive score. More details of neuropsychological assessments are described in Table S1 and the Methods section. Overall, participants showed decline in global cognition (Fig. S1).

**Table 1.** Demographic information for the cognitive reserve and reference ability neural network (CogRes/RANN) longitudinal study and the Alzheimer’s Disease Neuroimaging Initiative (ADNI) study datasets. The ADNI study dataset was used as an external dataset for generalizability validation.

| Dataset | Number of participants | Baseline age (SD) | Sex | Education (SD), years | Time (SD) from baseline to follow-up neuropsychological assessments, years |
| --- | --- | --- | --- | --- | --- |
| CogRes/RANN | 85 | 65.56 (3.35) | 42(F)/43(M) | 16.12 (2.22) | 4.51 (0.68) |
| ADNI | 72 | 73.69 (7.35) | 46(F)/26(M) | 16.83 (2.39) | ADNI-Mem and ADNI-GCog: 3.40 (1.18)<br>dMemory: 3.34 (1.17) |

### Association of global and regional Aβ with future cognitive decline

As linear statistical modeling is commonly used in the literatures for studying Aβ-cognition relationship^14,17,24,32^, we first performed a similar statistical association analysis before employing the cross-validated predictive modeling. The correlations between baseline Aβ pathology and prospective decline in each cognitive domain and in global cognition were assessed using Spearman partial correlation, controlling for age, sex, education, baseline cognition, and hemispherical mean cortical thickness. When using global SUVR of Aβ pathology, no significant relationships were observed (reasoning: r = 0.1321, p = 0.2369; memory: r = 0.1186, p = 0.2885; vocabulary: r = 0.2009, p = 0.0758; processing speed: r = −0.0526, p = 0.6387; global cognition: r = 0.0873, p = 0.4444). Next, we performed similar correlation analyses for each regional Aβ measure. The significance level was set as p < 0.05 with False Discovery Rate (FDR) multiple comparison correction for a total of 214 regions. We found significant relationships for the vocabulary cognitive domain, but not for reasoning, memory, processing speed, nor global cognition. Higher baseline RAP (PET regional SUVR values) in the left hemispherical extrastriate cortex (part of the central visual network) was significantly related to greater decline in vocabulary (r = −0.4044, FDR-corrected p < 0.0469; uncorrected p < 0.0002).

The nonsignificant results observed may be attributed to the use of a less sensitive non-parametric method, which was employed to accommodate the non-normal distribution of Aβ pathology^14^. Although weak but significant associations between Aβ deposition and future cognitive decline have been well established in the literature ^13–17^, the results of statistical association analyses can be influenced by the complexity of the statistical model and the sample size. Given that the Aβ-cognition relationship was evaluated using within-sample correlations, the results might not predict such relationship for the new individuals, and are not adequate to support predictive validity of early Aβ on cognition^33,34^. To improve generalizability in the findings of amyloid-cognition relationship, we next used cross-validated models.

### Baseline regional Aβ predicts subsequent cognitive decline in global cognition

As Aβ deposition has been reported to yield the largest effect size on global cognition^15^, we focused on this relationship. We employed a cross-validated predictive modeling approach, where a framework was adapted from connectome-based predictive modeling (CPM)^35–37^. Briefly, participants were randomly split into training and test partitions (Fig. 1a and 1b). Next, for the amyloid-cognition relationship, feature selection and model building were performed on the training set by Spearman correlation and linear modeling, respectively (Fig. 1a and 1c). Lastly, model predictive performance was evaluated on the correlation coefficient between predicted cognitive decline score and the actual cognitive change (cognition was assessed as residual value controlling covariates; Fig. 1b). More details are in the “Methods” section. As repeated k-fold cross-validation has proven to produce less biased estimation than the leave-one-out approach^38^, we used a 15-folds cross-validation modeling, and the random partitions of train and test sets were repeated 500 times to obtain a stable result (details of cross-validation fold number selection are in Supplementary materials). The predictive performance was evaluated as the median value across repeated iterations. First, we performed the cross-validated predictive modeling on RAP measures (regional SUVR values; Fig. 1a) to predict cognitive decline in global cognition. The same predictive modeling can be fitted to other pathology models such as NAP, incorporating either structural or functional connectome (see “pathology models” in Fig. 1a). For RAP measures, a median correlation coefficient R = 0.2013 was found between the predicted cognitive decline in global cognition and the actual values. To test its significance level, we generated an empirical null distribution by shufling the cognitive decline variable 500 times (the covariates were ordered the same each time as cognitive decline variable to keep correspondence). The p-value was calculated as the proportion of permutations that were greater than or equal to the actual correlation. We found that baseline RAP measures significantly predict subsequent cognitive decline in global cognition (p < 0.0459), indicating that the observed correlation coefficient R = 0.2013 is significantly higher than the amyloid-cognition relationship observed by chance.

**Fig. 1:**
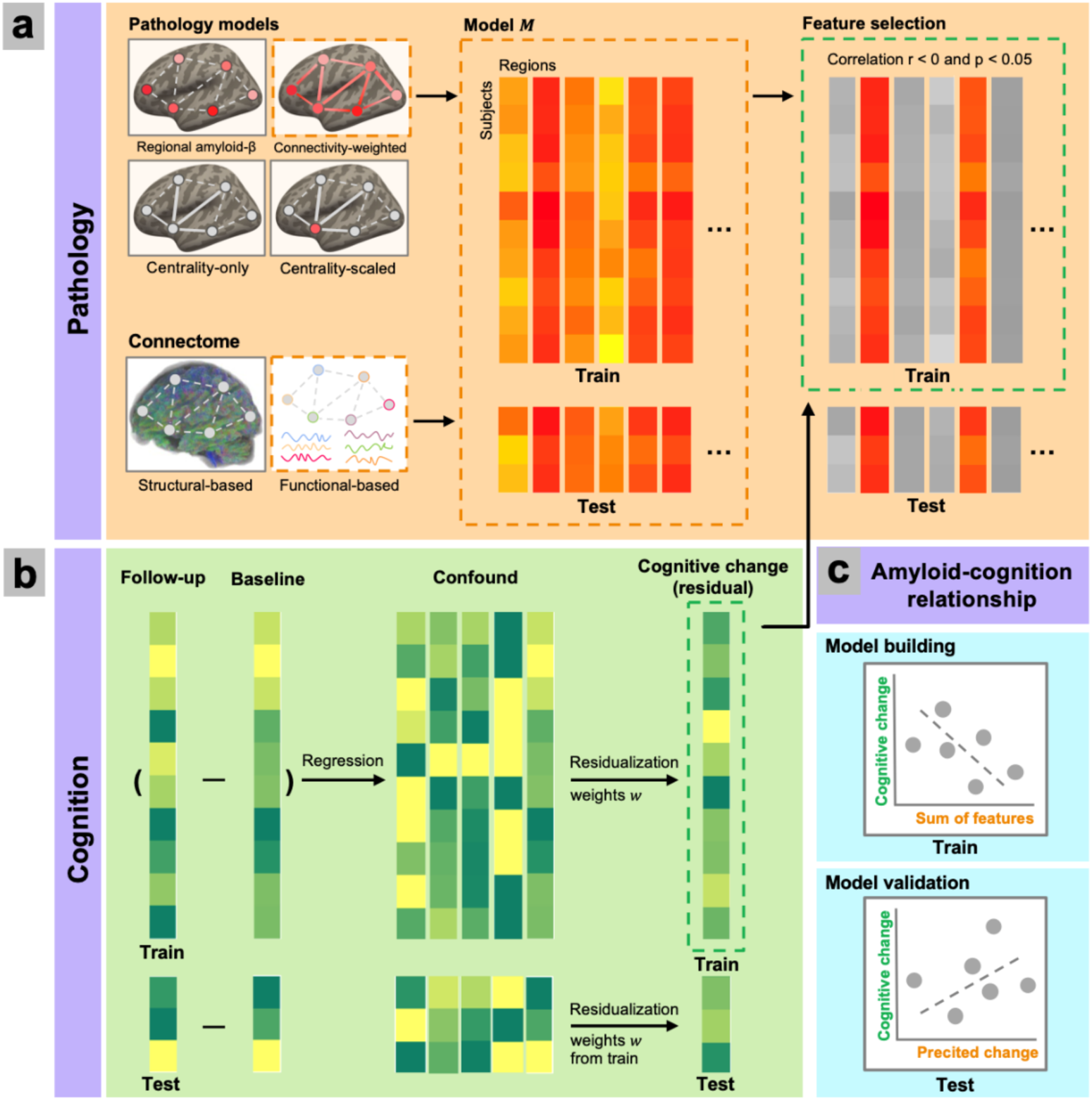
Cross-validated predictive modeling analysis overview. The figure illustrates an example of using connectivity-weighted network-level amyloid-β pathology (NAP) with functional-based connectome. **a** *Feature selection on the input of pathology models and connectome.* Regional amyloid-β pathology (RAP) or NAP (either connectivity-weighted or centrality-scaled) can be used as input. Individualized network connectomes can be defined based on either structural or functional imaging data. RAP measures do not need the input of connectome information, and the centrality-only model has only connectome information (serving as a benchmark for NAP measures). Since higher pathology is expected to associate with worse cognitive performance, a feature (i.e. regional measures of pathology) is selected if it shows a negative correlation with cognitive change residual score at an uncorrected p < 0.05 in the training set. **b** *Computation of longitudinal cognitive change scores.* Subjects are randomly split into a training set and test set. In the training set, we regress out covariate confounds from the longitudinal cognitive change. The covariate-related variability is also removed from the test set using the same weights. **c** *Regression of selected amyloid pathology features in the train set and prediction of cognitive decline in the test set.* The sum of selected features is regressed against cognitive change residual scores from the training sample to obtain model weights. The obtained weights are then applied to the features from test set to predict cogni7ve decline scores, which were then compared to actual scores to evaluate model predic7on performance.

### Connectome-based modeling of the Aβ improves prediction of subsequent cognitive decline

In addition to the regional Aβ deposition (Fig. 2a), we examined how the measures of Aβ deposition within individualized brain connectivity networks contribute to subsequent cognitive decline. Thus, we developed NAP scores by leveraging the connectivity profile of each region of interest (ROI) (Fig. 2b and 2c). As shown in Fig. 2e and 2h, the proposed connectivity-weighted NAP scores quantified the influence of Aβ deposition within the connected network associated with each ROI, where the NAP measure of each region quantifies not only the regional Aβ deposition in the region, but also Aβ in all other regions connected to this region, weighted by its connection strength to the region (More details are in the “Methods” section). A simpler version of NAP scores, i.e., centrality-scaled NAP, was also developed to quantify only the regional Aβ deposition in the region, and it is scaled by the centrality (nodal strength) of the region in the connectivity matrix (Fig. 2f and 2i). The rationale for such measures is that highly connected regions are crucial for cognitive functioning, and Aβ-related disruption of such regions may lead to more distributed effects on brain dynamics, and subsequently more severe cognitive decline^39^. Incorporating connectome information into pathology measures allowed us to identify network-based neuropathological patterns that might be predictive of cognitive decline. NAP measures could potentially uncover cognition-relevant pathological disconnection patterns. Specifically, we first obtained subject-specific structural and functional connectivity between each pair of ROIs through tractography and resting-state fMRI data, respectively (Fig. 2d and 2g; more details are in the “Methods” section). Then, we adopted the same cross-validated predictive modeling approach to evaluate and compare the performance of RAP and NAP measures, and additional models with only connectome centrality measures (no Aβ pathology information) were included as a benchmark for comparison.

**Fig. 2:**
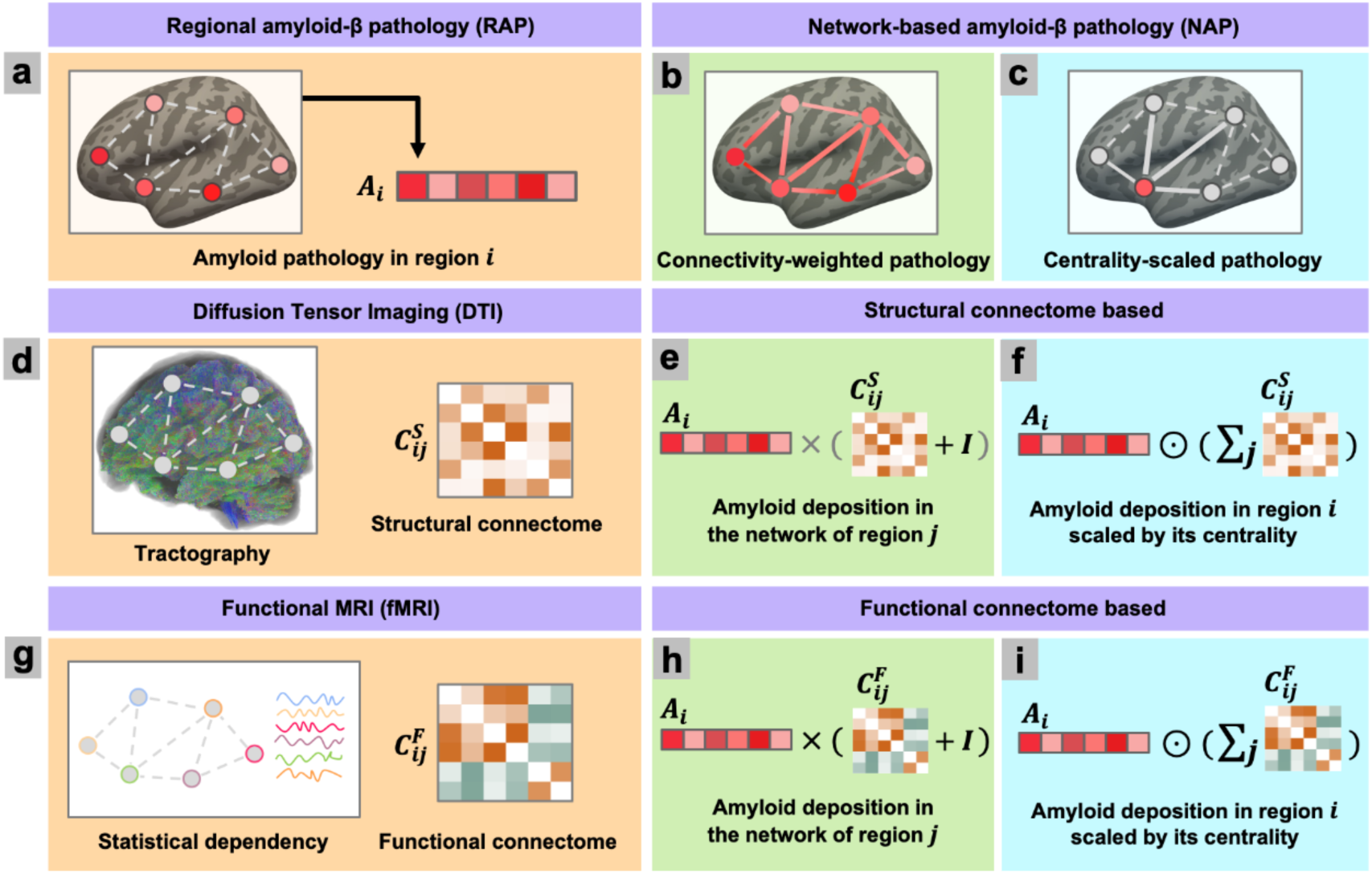
Regional amyloid-β pathology (RAP) and network-based amyloid-β pathology (NAP). In addition to RAP, the proposed NAP scores incorporate connectome information by using either a connectivity-weighted or centrality-scaled approach. Here, we show a six-region toy network. **a** Illustration of RAP deposition *A_i_* in region *i (i = 1, …, 6).* **b** Connectivity-weighted NAP score quantified the influence of Aβ deposition within the connected networks. **c** Centrality-scaled score quantifies the Aβ deposition in the region and scaled it by its centrality in the connectome. Network connectome information can be generated based on either structural connectome (SC) (**d**) or functional connectome (FC) (**g**). **d** SC is derived from tractography on diffusion tensor imaging data, where SC matrix was denoted as 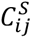. **e** SC based connectivity-weighted NAP score of region *j* quantifies both the Aβ in region *j* and a weighed sum of Aβ in all other regions based on connectivity values, where the whole process was denoted as matrix multiplication of 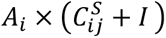. **f** SC based centrality-scaled NAP score of region *i* is quantified by a Hadamard product of RAP measure and connectivity centrality 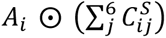. **g** FC is computed from statistical dependency on functional MRI data. **h, i** For the connectivity-weighted (**h**) and centrality-scaled pathology (**i**), FC based NAP scores are computed the same as SC based approach but with FC denoted as 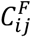.

To compare the predictive performance across pathology models, the train/test partitions for the cross-validation repetitions were kept the same for each model. Predictive performance was assessed by comparing predicted and actual cognitive decline scores with median correlation coefficient and ratio of average prediction error (denoted as *Ratio*, more details are in the “Methods” section). As shown in Fig. 3a and 3b, four different predictive models were built based on different network-based measures (connectivity-weighted or centrality-scaled measures) of brain connectome and pathology and were all compared to the predictive model purely based on regional amyloid SUVR. The performance difference in correlation coefficient and the *Ratio* were reported. In addition, two predictive models were also built based on the connectome information only as well (centrality-only measures). To test significance level of predictive performance, the same permutation procedure was employed by shufling the cognitive decline variable. While RAP measures predicted cognitive decline with a median correlation coefficient R = 0.2013, which is significantly higher than the amyloid-cognition relationship observed by chance (p < 0.0459). Structural connectivity information seems to be predictive of cognitive decline by itself, with centrality-only measures showing significant performance (median R = 0.2297, permutation p < 0.014). However, the prediction performance was the highest for the NAP measures when combining the information from both RAP and structural connectome information. Structural connectivity-based NAP measures also significantly predicted cognitive decline (connectivity-weighted NAP: median R = 0.2682, permutation p < 0.008; centrality-scaled NAP: median R = 0.2661, permutation p < 0.012). However, functional connectivity information itself had a worse performance compared to RAP, with a non-significant performance using centrality-only measures (median R = −0.0385, permutation p > 0.4950). Incorporating functional connectome information into the NAP measures made the predictive performance worse than RAP (connectivity-weighted NAP: median R = −0.0765, permutation p > 0.5968; centrality-scaled NAP: median R = −0.0786, permutation p > 0.5848). Additionally, we further compared the performance of structural connectome-based NAP and RAP measures. As shown in Fig. 3a, the performance of NAP measures is significantly higher than that obtained by RAP measures, with a difference in correlation coefficients significantly higher than zero (connectivity-weighted NAP: p < 0.004; centrality-scaled NAP: p < 0). The performance of centrality-only measures is higher than that obtained by RAP measures, but the difference is not significant (p > 0.1580).

**Fig. 3:**
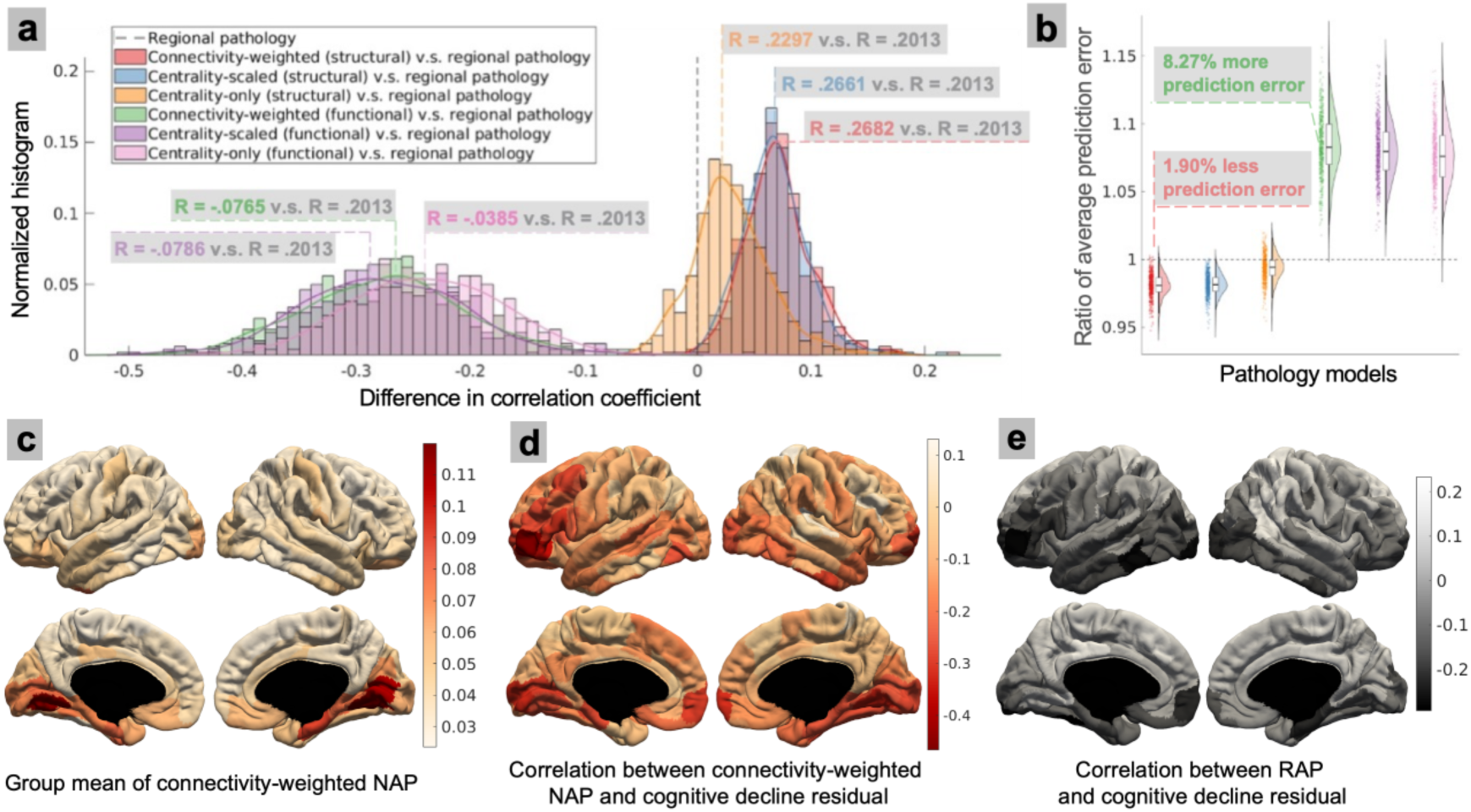
Performance of cognitive decline predicting performance using different pathology models across cross-validation repetitions. **a** Distributions of difference in correlation coefficients between predicted and actual cognitive decline scores across different pathology models. Prediction performance was compared across cross-validation repetitions with regional amyloid-β pathology (RAP) model as the reference. Incorporating structural connectome (SC) with connectivity-weighted network-based amyloid-β pathology (NAP) achieved the highest performance (median R = 0.2682), and outperformed (p < 0.004; test on difference in correlation coefficients against zero) the prediction performance of RAP measures (median R = 0.2013). **b** Comparison of ratio of average prediction error (*Ratio*) performance between pathology models. Compared to RAP measures, connectivity-weighted NAP measure of SC had *Ratio* = 1.90% less prediction error, whereas connectivity-weighted NAP measures of functional connectome (FC) had *Ratio* = 8.27% more prediction error. **c** Group averaged SC connectivity-weighted NAP scores. **d** Correlation coefficient between cognitive decline and SC connectivity-weighted NAP scores across cortical regions of interest (ROIs) (Spearman partial correlation, controlling for age, sex, education, baseline cognition, and hemispherical mean cortical thickness). **e** Correlation coefficient between cognitive decline and RAP scores across cortical ROIs. NAP scores have higher values in basal cortex regions (**c**). Individual variability in NAP scores in basal cortex regions and the left dorsolateral prefrontal cortex scales with cognitive decline (**d**), showing a stronger negative correlation with subsequent cognitive change compared to RAP scores (**e**).

In an additional analysis, to identify and visualize the brain ROIs contributing most to the observed relationships between pathology and cognition, we conducted a Spearman partial correlation analysis between the pathology features in each ROI and cognition change, controlling for age, sex, education, baseline cognition, and hemispheric mean cortical thickness. As shown in Fig. 3c, structural connectome-based, connectivity-weighted NAP scores had higher values in the basal cortical regions of the temporal and occipital lobes compared to other brain regions. These regions, along with the medial prefrontal cortex and left dorsolateral prefrontal cortex, exhibited a stronger negative correlation with longitudinal cognitive change than other areas of the brain (Fig. 3d). Furthermore, NAP scores showed a more pronounced negative correlation with longitudinal cognitive changes compared to RAP scores (Fig. 3e). Similar patterns were observed for the centrality-scaled NAP scores (Fig. S2).

While these results (Fig. 3a and 3b) were based on unthresholded connectome matrices, the overwhelming number of connections might make it difficult to extract meaningful information. Thus, we additionally tested whether keeping only meaningful connections above a proportional threshold will improve the performance of functional connectivity-based NAP measures. A proportional threshold approach was used as it has been shown to give more stable network measures than an absolute threshold approach^40^. As shown in Fig. S3a, a stricter threshold did improve the predictive performance with lower *Ratio* values compared to the performance of RAP. However, even the best performance model with a threshold of keeping the top 2% connections (connectivity-weighted NAP: median R = 0.1571; centrality-scaled NAP: median R = 0.0806) still performs worse than RAP measures (Fig. S3b).

### Forward application of identified neuropathological signatures predict future cognitive decline in an external dataset

Here, we employed a data-driven approach to identify neuropathological patterns of either regional or network-based Aβ deposition that maximally predicts future cognitive decline. To illustrate the results of feature selection in the cross-validated predictive modeling, Fig. 4 presents brain ROIs where pathology measures are predictive of future cognitive decline, by using RAP measures (Fig. 4a), connectivity-weighted NAP measures (Fig. 4b), or centrality-scaled NAP measures (Fig. 4c). These regions were identified if the features were selected for 95% of the iterations in cross-validation. As shown in Fig. 4, the Aβ neuropathological signatures identified regions located in basal portions of the frontal, temporal and occipital lobe, consistent with the early-stage Aβ deposition patterns reported in other PET and autopsy studies^26,41^. The exact region labels and location are reported in Table S2.

**Fig. 4:**
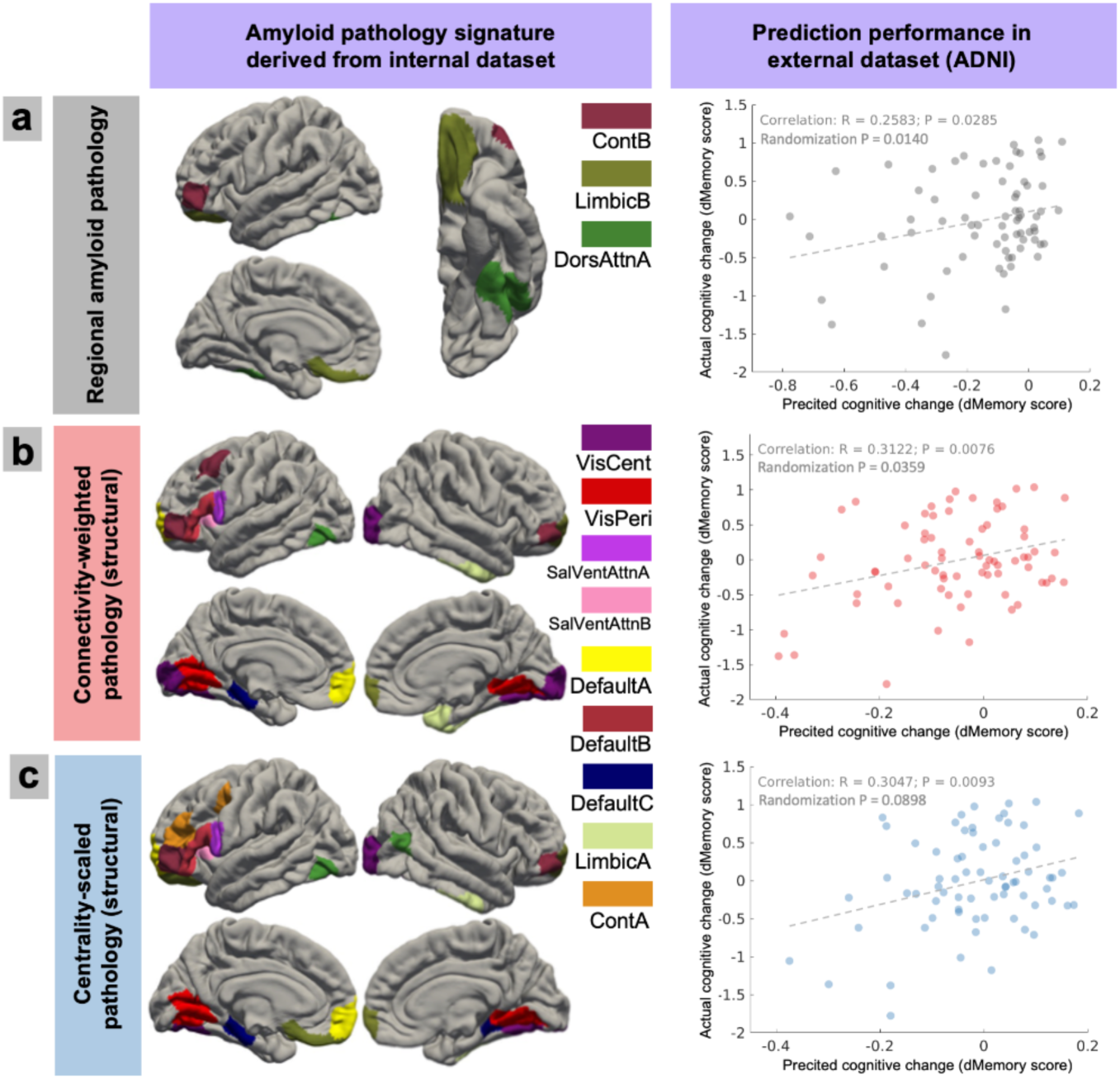
The amyloid-β (Aβ) neuropathological signatures of future cognitive decline and external validation. **a** Regions of interest (ROIs) whose regional Aβ neuropathology (RAP) measures were identified as features of future cognitive decline. This pattern of pathology signature was derived from the internal CogRes/RANN dataset. In external validation, features within the same ROIs pattern of ADNI participants were used to predict cognitive decline in delayed recall tests of memory (dMemory), and subsequently compared to the actual decline score. The predicted cognitive change significantly correlated with the actual score (R = 0.2583, p < 0.0285), and the relationship is significantly higher than the results obtained by random ROI selection (randomization p < 0.0140), demonstrating regional specificity. **b** Results of Aβ neuropathological signatures using connectivity-weighed network-based Aβ pathology (NAP) measures. **c** Results of Aβ neuropathological signatures using centrality-scaled NAP measures. The color indicates the network label of each ROI based on Schaefer atlas. Abbreviations: Cont, Cognitive control network; DorsAttn, Dorsal attention network; VisCent, Visual central network; VisPeri, Visual peripheral network; SalVentAttn, Salience/Ventral Attention network; Default, Default mode network.

We first validated the signature pattern in the same internal CogRes/RANN dataset. Features in the signature pattern were used to predict cognitive decline, and subsequently be related to the actual cognitive decline score (residual controlling age, sex, education, baseline cognition, and hemispheric mean cortical thickness) in global cognition, as well as in each cognitive domain. The predicted score significantly correlated with cognitive decline in both global cognition and episodic memory, but not in fluid reasoning, vocabulary, nor processing speed (Results shown in Table S3). Within the same dataset, the results indicate that the predictive validity of Aβ pathology in global cognition is mostly driven by its relationship to episodic memory. These are consistent with literatures showing the robust relationship between Aβ deposition and cognitive decline in episodic memory^13–17^. Next, we sought to validate the derived Aβ neuropathological signatures’ relationships to cognitive decline using an external dataset. We used Alzheimer’s Disease Neuroimaging Initiative (ADNI) dataset (adni.loni.usc.edu), and based on the findings using internal CogRes/RANN dataset, we focused on the cognitive decline in global cognition and episodic memory. Three cognition scores were extracted: 1) ADNI memory score (ADNI-Mem)^42^; 2) ADNI global cognition score (ADNI-GCog; summarized score of memory, executive functioning, and language)^43^; 3) Delayed recall tests of memory (dMemory; summarized score of logical memory delayed total recall and delayed word recall). More details of the cognition scores in ADNI dataset are described in the “Methods” section, and the longitudinal change in the ADNI cognition scores are illustrated in Fig. S4.

To test generalizability of the Aβ neuropathological signature, PET and MRI imaging data from the ADNI were preprocessed, and structural connectome and pathological measures were computed in the same way as the CogRes/RANN dataset (more details in the “Methods” section). The sum of pathology features in the signature pattern were used to predict cognitive decline in ADNI cognition scores (residual score controlling for covariates; see “Methods” section for more details). The predicted scores significantly correlated with the actual cognitive decline scores in dMemory (RAP: R = 0.2583, p < 0.0285; connectivity-weighted NAP: R = 0.3122, p < 0.0076; centrality-scaled NAP: R = 0.3047, p < 0.0093; Fig. 4), but not in ADNI-GCog (RAP: R = 0.1894, p < 0.1136; connectivity-weighted NAP: R = 0.2251, p < 0.0591; centrality-scaled NAP: R = 0.1925, p < 0.1078) and ADNI-Mem (RAP: R = 0.1456, p < 0.2257; connectivity-weighted NAP: R = 0.1306, p < 0.2776; centrality-scaled NAP: R = 0.1132, p < 0.3472).

Besides predictive performance on cognitive decline, we additionally tested if the observed relationship is specific to these particular sets of regions in the amyloid-β neuropathological signature, i.e. regional specificity. To do this, we performed a control analysis involving random selection of a set of ROIs, ensuring that the number of ROIs matched those selected in the Aβ neuropathological signature. Then, features in this random ROI set were used to predict cognitive decline, and the correlation coefficient with the actual cognitive decline score was recorded. By repeating this random selection process 500 times, we generated a distribution of correlation values from these random ROI selections. As shown in Table 2, when testing RAP and connectivity-weighted NAP measures, the results demonstrated that the Aβ neuropathological signature significantly outperformed the randomly selected ROIs in predicting ADNI-GCog (RAP: p < 0.0379; connectivity-weighted NAP: p < 0.0479) and dMemory (RAP: p < 0.0140; connectivity-weighted NAP: p < 0.0359; Fig. 4a and 4b), but only showed a trend in ADNI-Mem (RAP: p > 0.1836; connectivity-weighted NAP: p > 0.0619). However, centrality-scaled NAP measures did not show significant regional specificity in any of the cognition test (ADNI-Gcog: p > 0.5988; ADNI-Mem: p > 0.1856; dMemory: p > 0.0898; Fig. 4c). Together the results demonstrated that the Aβ neuropathological signature generalizes to an external dataset for predicting cognitive decline, and patterns derived from RAP and connectivity-weighted NAP demonstrated regional specificity.

**Table 2.** Model performance of amyloid-β neuropathological signatures on ADNI external dataset validation (* p < 0.05, p value in the regional-specificity control analysis was reported here instead of correlation p value). ADNI global cognition includes memory, executive function, and language cognition scores. dMemory (delayed memory recall) score includes logical memory delayed recall (LMDR) and delayed word recall from the Alzheimer’s Disease Assessment Scale–Cognitive Subscale (DWR-ADASc).

| Cognitive decline measures | Regional amyloid-β (Aβ) pathology | Connectivity-weighted network Aβ pathology | Centrality-scaled network Aβ pathology |
| --- | --- | --- | --- |
| ADNI global cognition score | $r = 0.1894$; $p = 0.0379$ (*) | $r = 0.2251$; $p = 0.0479$ (*) | $r = 0.1925$; $p = 0.5988$ |
| ADNI memory score | $r = 0.1456$; $p = 0.1836$ | $r = 0.1306$; $p = 0.0619$ | $r = 0.1132$; $p = 0.1856$ |
| dMemory score | $r = 0.2583$; $p = 0.0140$ (*) | $r = 0.3122$; $p = 0.0359$ (*) | $r = 0.3047$; $p = 0.0898$ |

Lastly, we evaluated whether the proposed NAP scores are more sensitive to cognitive decline than RAP scores when not using a cross-validation model, which may perform better due to optimized data-driven feature selection. We performed a Spearman partial correlation analysis between pathology features in each ROI and longitudinal cognitive changes, controlling for age, sex, education, baseline cognition, and mean cortical thickness of each hemisphere. As shown in Fig. 5, both NAP scores exhibited a significantly stronger negative correlation with cognitive changes compared to RAP scores in both the CogRes/RANN and ADNI datasets, where cognition was assessed using global cognition and dMemory, respectively. Network-level correlation results, based on networks defined by the Schaefer atlas, are provided in Fig. S5. In addition to regional specificity, the results demonstrated higher sensitivity of the proposed NAP scores to cognitive decline compared to RAP scores.

**Fig. 5:**
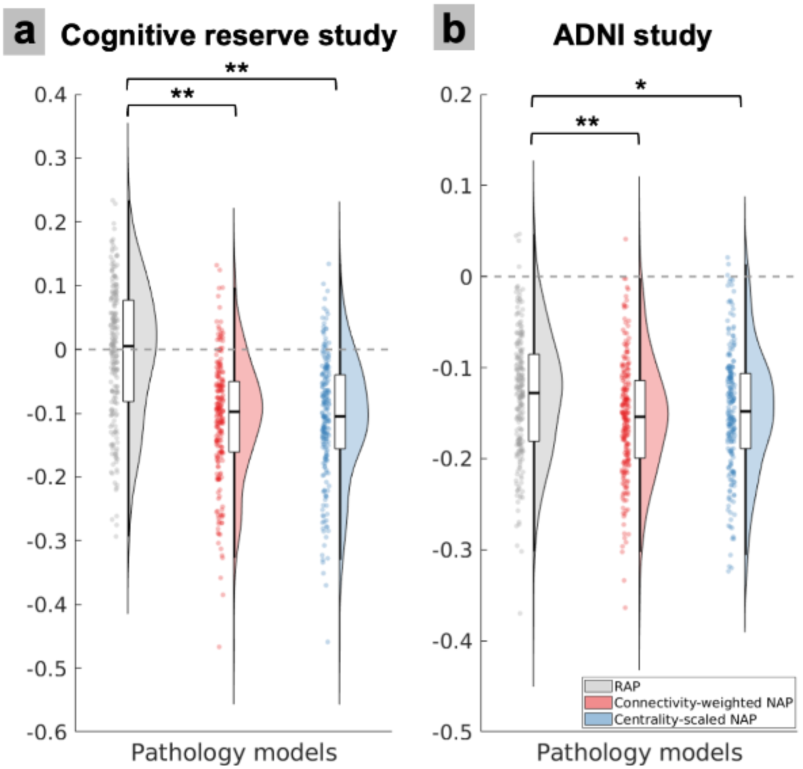
Correlation results in the relationship between pathology features in each region of interest (ROI) and longitudinal cognitive decline. **a** For participants in the cognitive reserve and reference ability neural network (CogRes/RANN) longitudinal study, we performed Spearman partial correlation analysis between pathology features in each ROI and longitudinal cognitive changes, controlling for age, sex, education, baseline cognition, and mean cortical thickness of each hemisphere. Pathology features were characterized by either regional amyloid-β pathology (RAP) or network-based amyloid-β pathology (NAP). **b** Same approach was performed for participants in the ADNI study. Compared to RAP measures, NAP scores demonstrated a significantly stronger negative relationship with cognition change in both studies. Cognition was assessed using global cognition in CogRes/RANN study and dMemory in ADNI study. (** p < 0.01; * p < 0.05; two-sample Student’s t-test)

## Discussion

Aβ deposition in the brain is an important hallmark of AD, sometimes occurring decades before the onset of cognitive impairment. However, the relationship between early deposition of Aβ and future cognitive decline is still unclear, especially since effective metric of brain Aβ that is predictive of future cognition is missing. Here, we aimed to establish a cross-validated predictive model to examine the proposed network-based Aβ pathology measures. A comprehensive examination was employed to evaluate predictive validity and external generalization. Our results demonstrated several important findings. First, early Aβ regional pathology significantly predicts future cognitive decline in global cognition. Second, incorporating structural connectome, but not functional connectome, information into pathology measures improves such predictive performance. Lastly, the Aβ neuropathological signatures derived from our internal dataset generalize to external dataset validation, i.e. features in such patterns were indicative of future cognitive decline, and the signature patterns demonstrated significant regional specificity.

Current correlation-based approaches for studying the amyloid-cognition relationship often fall short in providing informative insights for future studies, primarily due to the nature of their within-sample validation, which limits the generalizability of findings. In this study, we adapted a cross-validated predictive modeling approach, based on the CPM framework, to test the predictive validity of early Aβ deposition measures on future cognitive decline, where the performance was validated on the hold-out samples during cross-validation. Cross-validated predictive models, such as CPM, have been widely used to investigate brain-behavior relationships across various domains. For example, such models have been employed to study the relationship between resting-state functional connectivity and attention^35^, AD-related cognitive deficits^44^, task-based functional connectivity and creative ability^45^, cognitive reserve^46^, aggressive behavior^47^, as well as structural network lesioning with cognitive control^37^ and stroke-related motor impairments^48^. However, such approaches have not been commonly used to study neuropathology and cognitive aging. In this study, we demonstrated, using the cross-validated predictive modeling, that early Aβ regional pathology significantly predicts future cognitive decline in global cognition. Although previous research has established associations between Aβ deposition and cognitive decline in specific domains such as processing speed ^14^, executive function^49^, vocabulary and language^14,17^, and visuospatial function^15,50^, our study focused on global cognition because it is consistently reported to be associated with early Aβ deposition^14,15,50–52^, and demonstrated a stronger effect size compared to cognitions in individual domains^15^. Consistent with the literature, we did not observe significant relationships when models were trained on cognitive decline within cognitive domains. The stronger effect of Aβ on global cognition may reflect the downstream impact of Aβ on subsequent neuropathological processes, potentially contributing to a more pronounced relationship with clinical outcomes over time^18^. Our findings contribute to understanding regionally specific deposition of Aβ and its contributions to future cognitive decline.

It is well-established in the literature that early Aβ deposition exhibits a significant yet weak relationship with future cognitive decline. Given the multifaceted nature of AD, it is crucial to revisit the amyloid-cognition relationship by considering more comprehensive factors. Current studies often overlook the potential contribution of network-level metrics of neuropathology to subsequent cognitive decline. However, network-based approaches are important because neurological symptoms are increasingly understood to correlate more closely with dysfunction in brain networks rather than with deficits in specific locations. For instance, the lesion network mapping approach^53^ leverages the brain connectome to map common networks underlying lesions in different locations that manifest similar clinical symptoms. This method has been successfully applied to neuropsychiatric conditions such as amnesia^54^, depression^55^, and addiction remission^56^. Despite these advances, the network mapping of neuropathology in AD remains largely unexplored. Moreover, emerging evidence suggests that the spread of molecular neuropathology, as well as neurodegenerative damage, may follow large-scale brain networks^20,57^. Brain network organization could potentially mediate the relationship between regional neuropathology deposition and the heterogeneity observed in subsequent pathology accumulation and clinical symptoms^20^. Growing evidence shows that neuropathology distribution exhibits spatial heterogeneity and subtype structures. This was first observed in tau pathology^58^ and more recently in Aβ deposition, where distinct subtypes have been identified, such as frontal, parietal, and occipital predominant subtypes^29^ as well as temporal and cingulate predominant subtypes^25^. To relate network-based pathology and cognition, Luan et al. employed a lesion-network mapping approach to tau pathology, identifying distinct tau-lesion network maps associated with cognitive deficits in different cognitive domains^21^. Building upon such concepts, our study introduces novel NAP quantifications of Aβ deposition, utilizing subject-specific connectivity profiles to map networks associated with local Aβ deposition. Our approach is conceptually aligned with lesion network mapping^53^ and white-matter disconnection scores^37,48^, but we propose a more continuous and personalized metric that incorporates current understanding of the neuropathology underlying AD. For example, in our connectivity-weighted NAP measures, we use personalized connection strength as weights, reflecting the notion that stronger connections may facilitate faster pathology spread^20^. Additionally, our centrality-scaled measure is based on the idea that the pathology deposition in highly connected brain regions (regions of high topological centrality) may lead to more severe cognitive decline due to their widespread effects on brain dynamics^39^. It was also built upon the observations in the literature that Aβ pathology preferentially affects high centrality hub regions, demonstrating selective vulnerability in hub regions^59^. Centrality-based pathology measures, such as the tau hub ratio^60^, have been proposed to assess tau pathology and have been associated with cognitive decline. Our proposed centrality-scaled NAP measure builds on this prior work by utilizing subject-specific connectomes, offering a more individualized approach compared to group-average connectome methods. Our study provides the first characterization of network-level Aβ pathology and demonstrated that incorporating network connectome information into pathology measures enhances the predictive performance for subsequent cognitive decline compared to regional measures.

Our results highlighted the mediating role of individual connectome between neuropathology and cognition, which is important for early detection of AD-related cognitive deficits at the individual level. However, it remains unclear which attributes of the interplay between brain network architecture and neuropathology are responsible for the increased predictive performance. Potential candidate mechanisms are: 1) Individual connectomes capture coordinated regions that are selectively vulnerable to Aβ pathology. It has been shown that specific neurons, brain regions and brain networks exhibit selective vulnerability to neuropathology^8,61–64^. Highly connected hub regions with higher levels of neuronal activity and metabolic demand are vulnerable to neuropathology^59,65,66^. The proposed centrality-scaled NAP score, considering nodal centrality, may reflect networks with vulnerable hubs for the given subject, where severe AD-related symptoms might develop if Aβ disrupts highly functioning vulnerable hubs. 2) Individual connectomes are indicative of future Aβ spread and accumulation. Recent evidence suggested that many neurodegenerative disease-related pathological proteins (e.g. Aβ, tau, and α-synuclein) exhibit intercellular transmission^67^. This suggests that the early deposition of Aβ can potentially spread trans-neuronally across large-scale brain networks^65,68–70^. Thus, the selective vulnerability might be a result of the spreading between anatomically connected brain regions^67^. Some studies have shown that neuropathology progression along the structural connectome can be modeled to predict its future regional deposition^71^. The proposed connectivity-weighted NAP score, considering nodal connectivity profile, may suggest future Aβ spread based on its baseline deposition, where future deposition more closely tracks cognitive decline in the future. 3) Individual connectomes reflect the resilience of brain networks to neuropathology. Recent studies have found that topological characteristics of brain network resilience are related to age and cognition^72^. Incorporating connectome information into the pathology measures might account for individual resilience factors, and explain more variance in the cognition^18,73,74^.

Incorporating structural connectome into NAP measure significantly improves prediction performance on future cognitive decline, however, this is not the case for functional connectome-based NAP measures. One explanation is that functional connectivity reflects not only direct monosynaptic connections but also indirect polysynaptic connections^75^. Because functional connectivity quantifies statistical dependency between regional neural signals, the observed interactions between two regions might be mediated by other regions^76^. Trans-synaptic propagation of misfolded proteins might be shaped more by the structural than functional connections. Another possible reason is that the complex interplay between Aβ deposition and brain fMRI signals is often reported and not well understood. Some studies suggest that Aβ deposition leads to increased neural activity, and this hyperactivity may eventually lead to more rapid secretion of amyloid, forming a vicious cycle of Aβ accumulation and abnormal brain function^77^. The proposed simple NAP approach is not well suited to capture such complex interplay between fMRI signal and neuropathology. The present work may inform future studies on the complex interplay between brain network architecture, neuropathology, and cognition.

The neuropathological signature patterns identified in our study encompass regions within the default mode, cognitive control, visual, limbic, and salience networks, including the visual areas, temporal-occipital junction, temporal pole, insula, medial prefrontal cortex, lateral prefrontal cortex, orbitofrontal cortex, and parahippocampus. These regions are consistent with previous findings on early-stage Aβ deposition, where the initial deposition was observed in the basal portions of the frontal, temporal, and occipital lobes^26,78^. While studies reported early Aβ presence in basal visual areas^26,79^, other studies suggested amyloid reaches the visual cortex at later stages^7,41^. This inconsistency might be explained by recent work that identified an occipital subtype of Aβ accumulation, characterized by early abnormalities in the occipital cortex and a higher number of participants with dementia in such subtype compared to other subtypes^29^. Aβ deposition in the occipital cortex was often overlooked, whereas its deposition within occipital cortex has already been associated with cerebral amyloid angiopathy^80^ and Lewy body disease^81^. Recent studies also demonstrated neurophysiological measures during a visual processing task are sensitive to amyloid pathology in early-stage AD^82^. Our findings further support the notion that early Aβ deposition in the occipital lobe provides prognostic information for future cognitive decline. However, more studies are needed to investigate the relationship between Aβ occipital deposition and cognition. Moreover, the Aβ neuropathological signature derived from our internal dataset serves as a generalizable predictor of cognitive decline in the external dataset, demonstrating consistency relationship with cognition across datasets as well as regional specificity. However, the predictive power of our model is somewhat attenuated in the ADNI dataset, likely due to variations in PET radiotracers and diffusion MRI acquisition parameters that impact the structural connectome^83^.

This cross-validated predictive framework can be extended to other neuropathological and neurodegenerative markers, such as tau, α-synuclein, cortical atrophy, longitudinal Aβ accumulation^84^, or a composite measure of ATN (amyloid, tau, neurodegeneration) features^85^. Corresponding NAP scores for each measure can be computed to quantify its network-level effects and test their predictive performance on cognition. Additionally, future research should explore how cognitive reserve and resilience factors might mediate the relationship between Aβ accumulation and cognitive decline^18,74^. For example, increased brain activity and connectivity have often been observed alongside Aβ deposition^8^, potentially promoting cognitive reserve and helping to maintain cognitive function^86^. The high prediction performance of the proposed NAP might be attributed to its indication of future network propagation or network disruption of vulnerable region due to initial Aβ deposition and individualized connectome. This framework demonstrated superior predictive performance for future cognitive decline compared to regional Aβ pathology alone. However, these hypotheses on the underlying mechanisms need to be tested in future studies.

The current study has several limitations. First, the proposed framework needs to be validated with larger sample sizes to further ensure its robustness and generalizability. Second, since Aβ deposition is considered an early event in the neuropathological cascade of AD, we focused on the predictive validity of baseline Aβ deposition for future cognitive decline. However, the impact of Aβ on cognition is more complex, involving interactions with multiple factors that jointly affect cognitive outcomes. Specifically, in this study, we did not account for tau deposition or genomic information, such as APOE ε4 status, which may mediate the observed effects of Aβ on cognition. For example, evidence showed that high Aβ levels are associated with tau accumulation in the medial temporal lobe^4^, and Aβ and tau have interactive effects on cognitive decline^87^. Additionally, Aβ interacts with APOE ε4 to promote cognitive decline in cognitively normal older adults^88^, but APOE ε4 status was not included in our analysis. Aβ also interacts closely with neurodegenerative factors, such as hippocampal volume, glucose metabolism^89^, and cortical atrophy^90^. While we included hemispheric mean cortical thickness as a confounder, future studies should examine regional atrophy and its role in the Aβ-cognition relationship. Finally, future studies with longitudinal assessment of Aβ accumulation rates could provide additional insights into Aβ pathology^84^, however, recent evidence suggests that longitudinal measures may not significantly enhance the detection of cognitive decline compared to a single Aβ PET scan session^52^. In general, future research should comprehensively evaluate the potential factors that interact with Aβ pathology to better understand its relationship with future cognitive decline. Such a multimodal approach would be well-suited to capture the multifaceted nature of AD. Here, the proposed network-based approach and predictive modeling framework can be easily adapted to include other neuropathologies. And it would be intriguing to investigate an optimal combination of AD-related features, such as a composite score from ATN measures. These could be used to build and validate network-based predictive models for AD-related symptoms, similar to the framework proposed in this study, thereby promoting early detection of individuals on the AD continuum.

In summary, we developed a cross-validated predictive model to assess Aβ pathology measures for predicting future cognitive decline in cognitively normal older adults. Our results demonstrated that early regional Aβ pathology is a significant predictor of global cognitive decline, with predictive accuracy further enhanced by incorporating individualized structural connectome data as network-based Aβ pathology scores. The identified neuropathological signature pattern is indicative of future cognitive decline and demonstrates generalizability to external datasets. This network-based approach and predictive modeling complement existing pathology-based methods for early detection of individuals at risk of developing AD-related symptoms and can be extended to other neuropathological markers. The framework supports future neurodegenerative disease research that considers individual brain network connectivity when evaluating neuropathology and cognition.

## Methods

### Participants and study design in CogRes/RANN study

Participants were recruited through our ongoing CogRes/RANN longitudinal study^30^. All participants were recruited using random market mailing approach and were screened for basic inclusion criteria (i.e. right-handed, native English-speaking, no severe medical or psychiatric conditions, no head injuries, no hearing or vision impairments, and no other issues that could interfere with MRI acquisition or cognition). The experimental design of our study and the recruitment process were approved by the Internal Review Board of the College of Physicians and Surgeons of Columbia University. All participants have provided informed consent to participate in the study, and written consent was obtained from the participants. In this study, we included subjects who had: (1) participated baseline MRI acuiqistions and Aβ PET experiments, (2) demographic information available at baseline, and (3) finished neuropsychological testing at both baseline and follow-up sessions. Data were available for eighty-five subjects, with baseline age ranged from 56 to 71 (65.56 ± 3.35, mean ± SD) years, 42 females and 43 males, education ranged from 12 to 20 (16.12 ± 2.22, mean ± SD) years. To perform cross-validated predictive model analyses with NAP, 8 additional participants were excluded due to: 1) 3 subjects with functional connectivity results not passing quality control (QC); 2) 5 subjects with no baseline diffusion MRI data, with seventy-seven subjects left.

### Participants and study design in ADNI study

Data from the ADNI phase 3 (ADNI-3) database (adni.loni.usc.edu) were used in this study as an external validation dataset. The ADNI study received approval from the Institutional Review Boards of all participating institutions. And informed written consent was obtained from all participants at each site for participating in the ADNI repository. In this study, we included a sample of 159 cognitively normal older adults from a previous study^91^. In ADNI, cognitively normal subjects were identified as non-depressed, non-MCI, non-demented, with Mini-Mental State Examination (MMSE) scores of 24-30 and clinical dementia rating (CDR) scores close to zero. Additionally, we excluded 12 participants due to progression to MCI in the follow-up, 44 participants with no diffusion data acquisition or missing of information in diffusion data, 5 participants failed with structural connectivity QC, 7 participants failed with PET Aβ analysis, 18 participants without follow-up cognition available, 1 participant without baseline education available. Totally, in the external validation dataset, we had 72 participants, with baseline age ranged from 56.0 to 91.4 (73.69 ± 7.35, mean ± SD) years, 46 females and 26 males, education ranged from 12 to 20 (16.83 ± 2.39, mean ± SD) years. When testing the Aβ-cognition relationship with ADNI memory score and ADNI global cognition score, one additional participant was excluded due to follow-up ADNI memory score not available.

### Cognitive behavioral measures in CogRes/RANN study

A standardized battery of neuropsychological assessments was administered to assess cognition. In this study, we used a subset of the test results to assess cognition in four domains: 1) episodic memory (Selective Reminding Task (SRT) immediate recall, delayed recall, and delayed recognition^92^); 2) vocabulary (Wechsler Adult Intelligence Scale (WAIS-III) Vocabulary^93^, Wechsler Test of Adult Reading (WTAR)^94^, and American National Adult Reading Test (AMNART)^93^); 3) processing speed (WAIS-III Digit Symbol^93^, Stroop Color Word Test^95^, and Trail-Making Test versions A^96^), and 4) fluid reasoning (WAIS-III Matrix Reasoning^93^, WAIS-III Block Design^93^, and Trail-Making Test versions B^96^) (See the list of all the neuropsychological assessments employed in the CogRes/RANN study in Table S1). Based on the factor structure of these tests, cognitive composite scores for each cognitive domain were generated. To normalize the composite measures, z-scores were calculated within each domain, by subtracting the baseline full sample mean from each score and then dividing by the baseline full sample standard deviation. The average domain z-score for processing speed was reversed in sign, to ensure that higher values represented better cognitive performance. An average of the four z-scores of all four cognitive domains was calculated to yield a global cognition score. The baseline and follow-up neuropsychological assessments duration is 3 to 7 (4.51 ± 0.68, mean ± SD) years.

### Cognitive behavioral measures in ADNI study

Three ADNI cognition scores were assessed: 1) ADNI-Mem; 2) ADNI-GCog (summarized score of memory, executive functioning, and language); 3) dMemory. ADNI-Mem was assessed with modern psychometric approaches^42^, where Rey Auditory Verbal Learning Test (RAVLT, 2 versions), AD Assessment Schedule – Cognition (ADAS-Cog, 3 versions)^97^, MMSE, and Logical Memory data were analyzed, and a composite memory score was computed. ADNI executive functioning score (ADNI-EF) was derived from confirmatory factor analysis, and the final model included Category Fluency—animals, Category Fluency—vegetables, Trails A and B, Digit span backwards, WAIS-R Digit Symbol Substitution, and 5 Clock Drawing items (circle, symbol, numbers, hands, time). More details were described in^98^. ADNI language score (ADNI-Lan) was derived from MMSE (object naming, sentence repetition, sentence reading and writing, and following a three-step command), ADAS-Cog (following commands, object naming, and ideational praxis)^97^, ADNI administered Montreal Cognitive Assessment (MoCA) (phonemic fluency and sentence repetition)^99^. More details were described in^43^. These three ADNI composite scores were averaged to compute an assessment of global cognition (ADNI-GCog). To further assess episodic memory, we additionally derived a summarized score dMemory using delayed recall tests. Specifically, logical memory delayed total recall from the neuropsychological battery was converted to z-scores based on the baseline full sample mean and standard deviation from all cognitively normal older adults in ADNI-3. The same process was performed for the delayed word recall (item 4 of ADAS-Cog) to derive the z-scores. These two scores were averaged to obtain the dMemory score for assessing episodic memory. Follow-up cognition was chosen as the latest neuropsychological testing when multiple follow-up sessions were available. The baseline and follow-up neuropsychological assessments duration is 1 to 5.5 (3.40 ± 1.18, mean ± SD) years for ADNI-Mem and ADNI-GCog, and 1 to 5.5 (3.34 ± 1.17, mean ± SD) years for dMemory.

### Data acquisition and preprocessing in CogRes/RANN study

MR images were acquired using a 3 Tesla Philips Achieva Magnet scanner. An anatomical T1- weighted structural image was acquired using magnetization-prepared rapid gradient echo (MPRAGE) (TR/TE = 6.6/3.0 ms; flip angle = 8°; field of view (FOV) = 256 × 256 mm; matrix size = 256 × 256 voxels; voxel size = 1 × 1 × 1 mm; 165 axial slices). A neuroradiologist reviewed each participant’s MRI scan and confirmed the absence of clinically significant findings in all participants. Resting state fMRI scans were collected with a T2*-weighted echo-planar imaging (EPI) sequence (TR/TE = 2000/20 ms; flip angle = 72°; FOV = 224 × 224 mm; in-plane resolution = 112 × 112 voxels; slice thickness = 3 mm; 37 axial slices; either 150 or 285 volumes). Three dummy EPI volumes acquired at the beginning of each fMRI acquisition run were excluded before data preprocessing. Diffusion MR images were acquired with spin-echo echo-planar diffusion-weighted sequences. Participants’ data were acquired with a single shell sequence (TR/TE = 7647/69 ms, matrix size = 224 × 224 mm, voxel size = 2 × 2 × 2 mm, 75 axial slices, b = 800 s/mm2 with 56 gradient directions, and 1 volume with b = 0 s/mm2). Two runs of diffusion MR were performed for each subject, and data were merged before preprocessing.

Structural T1-weighted MR images were processed using the FreeSurfer pipeline v5.1^100^, which included segmentation of brain tissue, reconstruction of the cerebral cortex surface, and computation of cortical thickness. Hemispherical mean cortical thickness was computed as the average cortical thickness measures across 34 regions from the FreeSurfer Desikan-Killiany atlas^101^ for each hemisphere. Brain parcellation was performed based on the Local-Global Schaefer cortical parcellation atlas^102^ and FreeSurfer subcortical segmentation^103^. Specifically, 200 cortical ROIs were defined using the Schaefer atlas. The Schaefer atlas was aligned to each subject’s cortical surface through surface-based registration with FreeSurfer. Subsequently, the ROI surface areas were projected into volumetric space, resulting in the parcellation of the cerebral cortex into 200 ROIs. The Schaefer atlas grouped these 200 ROIs into 34 brain networks. Additionally, 14 subcortical ROIs were included from the FreeSurfer automatic segmentation^103^ (regions included bilateral thalamus, caudate, putamen, ventral striatum, globus pallidus, amygdala, and hippocampus). Totally, 214 ROIs were used with 200 cortical regions from the Schaefer atlas and 14 subcortical regions from FreeSurfer segmentation. Tissue segmentation and spatial registration were manually reviewed for quality control purposes^104^.

Participants underwent Aβ PET imaging with [^18^F]Florbetaben (donated by Piramal Pharma, Inc.) in a Siemens Biograph64 mCT/PET scanner (dynamic, 3D acquisition mode, 4 × 5-minute frames over 20 minutes). Brain PET image acquisition started 50 minutes after the bolus injection of 18F-florbetaben (10 mCi dose). An accompanying structural CT scan (in-plane resolution = 0.58 × 0.58 mm; slice thickness = 3 mm, FOV = 300 × 300 mm, and number of slices = 75) was acquired for attenuation correction purposes. PET data were reconstructed using the TrueX (HD-PET) algorithm and smoothed with a 2-mm Gaussian kernel with scatter correction.

### Data acquisition and preprocessing in ADNI study

In ADNI-3 study, MRI scanner protocols and parameters can be found in (https://adni.loni.usc.edu/data-samples/adni-data/neuroimaging/mri/mri-scanner-protocols/). Briefly, MR images were acquired with ADNI standardized pulse sequence at various sites using 3 Tesla scanners. Anatomical T1-weighted structural image was acquired using MPRAGE (TR/TE = 2300/2.96 ms; FOV = 240 × 256 mm; matrix size = 240 × 256 voxels; voxel size = 1 × 1 × 1 mm; 208 sagittal slices). Diffusion MR images were acquired with spin-echo echo-planar diffusion-weighted sequences. Most participants’ data were acquired with a single shell sequence (TR/TE = 7200/56 ms, matrix size = 232 × 232 mm, voxel size = 2 × 2 × 2 mm, 80 axial slices, b = 1000 s/mm2 with 48 gradient directions, and 7 volumes with b = 0 s/mm2). However, sequence parameters are different for different scanners, please see details in supplementary materials. To employ Aβ PET imaging, participants were administered with an intravenous catheter of [^18^F]Florbetapir (AV45, 10 mCi ± 10% dose). Then, a low-dose CT image for PET was acquired, followed by a brain PET image acquisition started 50 - 70 min after the injection (dynamic, 3D acquisition mode, 4 × 5-minute frames over 20 minutes). The same structural data preprocessing, as conducted on the CogRes/RANN study participants, were performed on ADNI data including FreeSurfer pipeline, brain parcellation with Schaefer atlas, subcortical ROIs delineation with FreeSurfer. However, a newer version of the FreeSurfer (v7.0) was used in ADNI data preprocessing.

### PET regional amyloid-β SUVR analysis

An in-house developed automatic quantification approach was used to analysis Aβ PET scans^105^. Briefly, four dynamic PET frames were aligned to the first frame using rigid-body registration and averaged to create a static PET image. This static PET image was then co-registered with the accompanying CT scan to generate a composite image. Then, each participant’s structural T1w image was registered to the composite PET-CT image (rigid-body registration with cost function as mutual information). ROIs and the cerebellar gray matter mask were transformed into the static PET image space. And regional PET data were extracted from these ROIs including 200 cortical and 14 subcortical regions. Lastly, standardized uptake value (SUV) was calculated for each ROI and was normalized to the value in the gray-matter cerebellum mask to derive SUVR. The same amyloid-β SUVR analysis was performed on ADNI PET imaging data.

### fMRI preprocessing and functional connectivity analysis

An in-house developed method was used to derive functional connectivity from resting-state fMRI data^106^. Firstly, slice timing correction was carried out with Fourier-space time-series phase-shifting by using FSL software package (version 6.0.4)^107^. Then, motion correction was performed using rigid-body registrations on all the volumes in reference to the first time point volume^108^. An fMRI reference image was created by averaging all the aligned EPI volumes. After that, frame-wise displacement (FWD) was calculated for each subject^109^, using the six motion parameters and the root mean square difference (RMSD) of the realigned fMRI signal between consecutive volumes. Scrubbing was performed to exclude fMRI contaminated volumes (if FWD exceeded 0.5 mm or RMSD surpassed 0.3%). Specifically, contaminated volumes were replaced with new ones generated through linear interpolation of adjacent volumes. Next, band-pass filtering was carried out with cut-off frequencies of 0.01 and 0.08 Hz, using the FSL software package with nonlinear high-pass and Gaussian linear low-pass. Finally, after motion-correction, scrubbing, and temporal filtering, we regressed out the FWD, RMSD, white matter signals from both hemispheres, and lateral ventricular signals from the preprocessed fMRI data. No global signal regression was applied. The averaged fMRI time-series signals were extracted from the 214 ROIs, including 200 cortical ROIs from the Schaefer atlas and 14 subcortical ROIs from the FreeSurfer segmentation. Then, functional connectivity matrices were calculated with Pearson’s correlation, and the correlation matrices were Fisher z-transformed to generate normalized connectivity matrices with the diagonal set to zeros. QC was performed based on^110^, by computing the variance of the mean signal within each ROI. Participants were excluded if at least one region had zero (or very low) variance.

### DTI preprocessing and structural connectivity analysis

Structural connectivity was generated following the procedures outlined in Basic and Advanced Tractography (BATMAN)^111^ and using the MRtrix3 software^112^. Firstly, diffusion MRI data were preprocessed with steps of denoising^113^, Gibb’s ringing artifact correction^114^, EPI geometric distortion correction^115^, eddy current and motion distortion correction^116^, bias field correction^117^, and brain mask estimation. Then, response functions for white matter, grey matter, and cerebrospinal fluid were estimated, followed by intensity normalization to correct global intensity differences. Whole-brain streamlines were generated through anatomically constrained tractography (ACT), creating 10 million streamlines per subject. The tractograms were then filtered using the spherical-deconvolution informed filtering of tracks (SIFT) to improve streamline density distribution. Finally, structural connectivity matrices were created using the predefined 214 cortical and subcortical ROIs extracted from the Schaefer atlas and FreeSurfer segmentation. The streamline count was normalized with regard to the inverse of the node volumes. The connectivity matrices were symmetrized, and the diagonal was set to zero. QC was performed based on^110^, specifically, we computed QC measures for connectome matrix density, number of connected components, and reconstructed streamlines. Participants were excluded if had more than one connected component, or either matrix density or number of streamlines fell outside three standard deviations from the group mean.

### Network-based pathology measures

The proposed connectivity-weighted NAP score was computed as the matrix product of regional Aβ deposition *A*_*i*_ in region *i = 1, …, N* (*N* regions of interest) and connectivity matrix *C*_*ij*_between region *i* and *j* (*j = 1, …, N*), denoted as: *A*_*i*_ × (*C*_*ij*_ + *I*). The connectivity-weighted (CW) NAP deposition at region *i* quantifies not only the regional Aβ deposition *A*_*i*_, but also Aβ deposition in all other regions *j* (*j = 1, …, N*) connected to region *i,* weighted by the strength of their structural or functional connectivity: 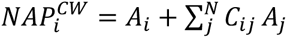. The proposed centrality-scaled (CS) NAP score was quantified by a Hadamard product of regional Aβ deposition *A*_*i*_ in region *i = 1, …, N* (*N* regions of interest) and the centrality of region *i* in the connectivity matrix *C*_*ij*_, denoted as: 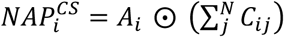. The centrality-scaled NAP score quantified regional Aβ deposition *A*_*i*_, and scaled it by its centrality regarding the whole connectivity matrix *C*_*ij*_. In both NAP scores, connectivity matrix can be generated either using structural diffusion data or functional MR data, denoted as 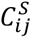 or 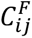, respectively.

### Cross-validated predictive modeling

Here, we adapted CPM to evaluate the Aβ-cognition relationship. Because leave-one-out approach has been shown to introduce biased estimates, repeated random splits of K-fold cross-validation modeling was chosen as a more stable approach^34,38^. We employed 500 repetitions to account for the impact of random train test data partitions, and the median prediction performance was reported. The cross-validated predictive modeling included four steps as shown in Fig. 1: 1) Participants splitting: subjects were randomly split into K-fold (K = 15 was chosen in this study, see Supplementary materials for details), including a training set and a test set. In the training set, as shown in Fig. 1b, we first regressed out covariates related variability (age, sex, education, corresponding baseline cognition, and hemispherical mean cortical thickness) from the longitudinal cognition change. This covariate-related variability was also removed from the test set using the same coefficient weights from the training set. 2) Feature selection: In the training set, the pathology measure (either regional or network-based amyloid) of each ROI was correlated with the longitudinal cognitive change (residual from the first step). Features/regions were selected showing a strong negative correlation with an uncorrected p value less than 0.05 (because elevated pathology deposition was expected to associate with severe cognitive decline). 3) Model training: In the training set, the sum of selected features was regressed against residual cognitive change to obtain model weights. 4) Model validation: In the test set, the sum of the features in the same selected ROIs and the model weights from step 3) were used to compute predicted cognitive change. In the loop of each hold-out fold, the same processes were performed, resulting in a predicted cognitive change value for each participant. Lastly, statistical analyses were performed to compare predicted and the actual cognitive change values in the collection of all hold-out samples. The whole process of cross-validation was repeated 500 times. An Aβ neuropathological signature pattern was derived for each of the pathology measures input. Regions in the pattern were identified if the regional features were selected 95% times during the repetition of cross-validation.

### Statistical analysis of predictive model performance

For each model, two metrics were computed to evaluate the predictive performance including Pearson’s correlation coefficient and average prediction error as 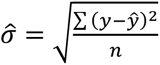, where *y* and 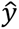 are the actual and predicted cognitive decline scores, respectively, and *n* denotes the sample size. To compare the predictive performance across pathology models, the train test partitions in the 500 times repetition were kept the same for each model. Each model was then compared to the performance of the RAP measure, which served as the reference model. We report the median correlation coefficient, difference in correlation coefficient to the reference model, and the ratio of average prediction error over the reference model. The *Ratio* quantifies the prediction error of each model compared to the RAP reference model and was computed as the average prediction error 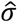 of the model divided by that of the reference model. Between each model and the RAP reference model, an empirical one-sided sign test was performed on the histogram of difference in correlation coefficient, and p values were reported.

### External generalizability analysis on ADNI dataset

As evidenced in our findings and existing literature^13–17^, Aβ-cognition relationship is consistently observed in the episodic memory domain and global cognition. Therefore, in the generalizability external validation, we tested the Aβ neuropathological signatures derived from internal CogRes/RANN dataset on the following neuropsychological assessments in the ADNI-3 dataset: 1) ADNI-Mem; 2) ADNI-GCog; 3) dMemory. Using the same approach as that used for the cross-validated predictive model analysis, residual of the longitudinal cognitive change (follow-up minus baseline) was computed by regressing out covariates including age, sex, education, hemispherical mean cortical thickness, and the corresponding baseline cognition. However, because ADNI-3 participants had a large variability in the time interval between baseline and follow-up cognition, this time interval variability was included as an additional covariate.

The same structural T1-weighted MR data analyses were applied to ADNI-3 dataset, including brain parcellation pipelines and cortical thickness estimation with FreeSurfer. However, a newer version of the FreeSurfer (v7.0) was used. Manual inspection and editing were conducted by a technician to ensure quality control on cortical surface reconstruction, and white and gray matter borders delineation. The diffusion MR data used for each participant were selected based on the closest time interval between data collection and the baseline neuropsychological assessments. Then, the same preprocessing and structural connectome procedures were performed to derive structural connectivity in the ADNI-3 cohort. However, the pipeline was slightly adjusted for three participants due to the use of a different acquisition sequence involving a multi-shell scheme.

For the external generalizability validation on the Aβ neuropathology signature, we extracted the features (RAP, connectivity-weighted NAP, or centrality-scaled NAP values) in the corresponding signature pattern for each pathology model. Then, the sum of features was used to predict cognitive decline, using the same weights estimated on the internal dataset. To evaluate the performance, Pearson’s correlation was calculated between the predicted values and the actual cognitive decline residual scores in each of the ADNI cognition (ADNI-Mem, ADNI-GCog, or dMemory). Additionally, we tested whether the observed relationship reflects general Aβ deposition in the brain, rather than being specific to the distinct signature patterns that are characteristic of future cognitive decline. To test such regional specificity, we conducted a control analysis where we randomly selected a set of ROIs, ensuring that the number of ROIs matched those identified in the Aβ neuropathological signature. Then, for each iteration, we used the features from this randomly selected set of ROIs to predict cognitive decline, and we recorded the correlation coefficient between the predicted values and the actual cognitive decline scores. This process was repeated 500 times, resulting in a distribution of the correlation coefficients. Lastly, the observed relationship in the signature ROIs was compared to this control analysis distribution, and the p values in the regional specificity randomization test were reported. The p-value was calculated as the proportion of randomizations that produced a correlation greater than or equal to the observed prediction correlation.

## Data and code availability

The CR/RANN datasets analyzed in the current study are available from the corresponding author on reasonable request. The code and sample data are available at https://github.com/hehengda/cross_validation_network_based_amyloid.git.

## Acknowledgments

This work was supported by the National Institutes of Health/National Institute on Aging (NIH/NIA; grant numbers R01 AG038465 and R01 AG026158)

## Author contributions

Design of research: H.H., C.H. and Y.S.; Methodology: H.H. and C.H.; Formal analysis: H.H.; Data collection: Y.S.; Writing original draft: H.H.; Writing, review, and editing: H.H., Q.R., Y.G., C.H. and Y.S.; Funding and supervision: C.H. and Y.S.;

