## Supplementary Information for "Network-based amyloid-β pathology predicts subsequent cognitive decline in cognitively normal older adults"

### Supplementary text

ADNI diffusion data were collected from different scanners with different sequence parameters as the following: 1) 45 participants' data were acquired from a Siemens Prisma scanner with a single-shell sequence of 42 participants (TR/TE = 7200/56 ms, matrix size = 232 × 232 mm, voxel size = 2 × 2 × 2 mm, 80 axial slices, b = 1000 s/mm2 with 48 gradient directions, and 7 volumes with b = 0 s/mm2) and a multi-shell sequence of 3 participants (TR/TE = 3300/71 ms, matrix size = 232 × 232 mm, voxel size = 2 × 2 × 2 mm, 80 axial slices, b = 500, 1000, 2000 s/mm2 with 112 total diffusion weighted directions); 2) 8 participants' data were acquired from a Siemens Skyra scanner (TR/TE = 9600/82 ms, matrix size = 232 × 232 mm, voxel size = 2 × 2 × 2 mm, 80 axial slices, b = 1000 s/mm2 with 48 gradient directions, and 7 volumes with b = 0 s/mm2); 3) 10 participants' data were acquired from a Siemens Verio scanner (TR/TE = 16,700/105 ms, matrix size = 128 × 128, 80 axial slices, voxel size= 2.67 × 2.67 × 2 mm3, b = 1000s/mm2 for 30 gradient directions and one volume with b = 0 s/mm2); 4) 2 participants' data were acquired from a Siemens Trio scanner (TR/TE = 12,400/95 ms, matrix size = 232 × 232 mm, 80 axial slices, voxel size= 2 × 2 × 2 mm3, b = 1000s/mm2 for 30 gradient directions and one volume with b = 0 s/mm2); 5) 5 participants' data were acquired from a Philips Achieva scanner (TR/TE = 10,013/87 ms, matrix size = 256 × 256 mm, 80 axial slices, voxel size= 2 × 2 × 2 mm, b = 1000s/mm2 for 32 gradient directions and one volume with b = 0 s/mm2); 6) 1 participant's data were acquired from a Philips Ingenia scanner (TR/TE = 10,861/100 ms, matrix size = 256 × 256 mm, 80 axial slices, voxel size= 2 × 2 × 2 mm, b = 1000s/mm2 for 32 gradient directions and one volume with b = 0 s/mm2); 7) 1 participant's data were acquired from a Siemens BioGraph mMR scanner (TR/TE = 12,400/95 ms, matrix size = 232 × 232 mm, 80 axial slices, voxel size= 2 × 2 × 2 mm3, b = 1000s/mm2 for 30 gradient directions and one volume with b = 0 s/mm2)

We tested the predictive performance of the cross-validation model using regional amyloid-β pathology to predict global cognition decline, with the fold number K range in 5, 10, 15, and 20. The results showed that the model yielded the best predictive performance when K = 15 (median correlation coefficient R; 5 folds: R = 0.1985; 10 folds: R = 0.2010; 15 folds: R = 0.2013; 20 folds: R = 0.2004).

To test the relationship between baseline regional amyloid deposition and subsequent longitudinal cognitive decline, the predictive modeling approach was additionally employed for cognitive decline values in each separate cognitive domain (episodic memory, vocabulary, processing speed, and fluid reasoning). While baseline amyloid-β pathology yielded significant results in predicting cognitive decline in global cognition, no significant results were found for each separate cognitive domain (episodic memory: median R = 0.0732, permutation p > 0.2854; vocabulary: median R = 0.0848, permutation p > 0.2136, processing speed: median R = -0.0953, permutation p > 0.5868, and fluid reasoning: median R = 0.1067, permutation p > 0.1557). These results align with previous findings suggesting that Aβ has a more pronounced relationship with global cognition compared to specific cognitive domains[1].

**Fig. S1.** Longitudinal change in the cognitive reserve and reference ability neural network (CogRes/RANN) longitudinal study cognition scores.

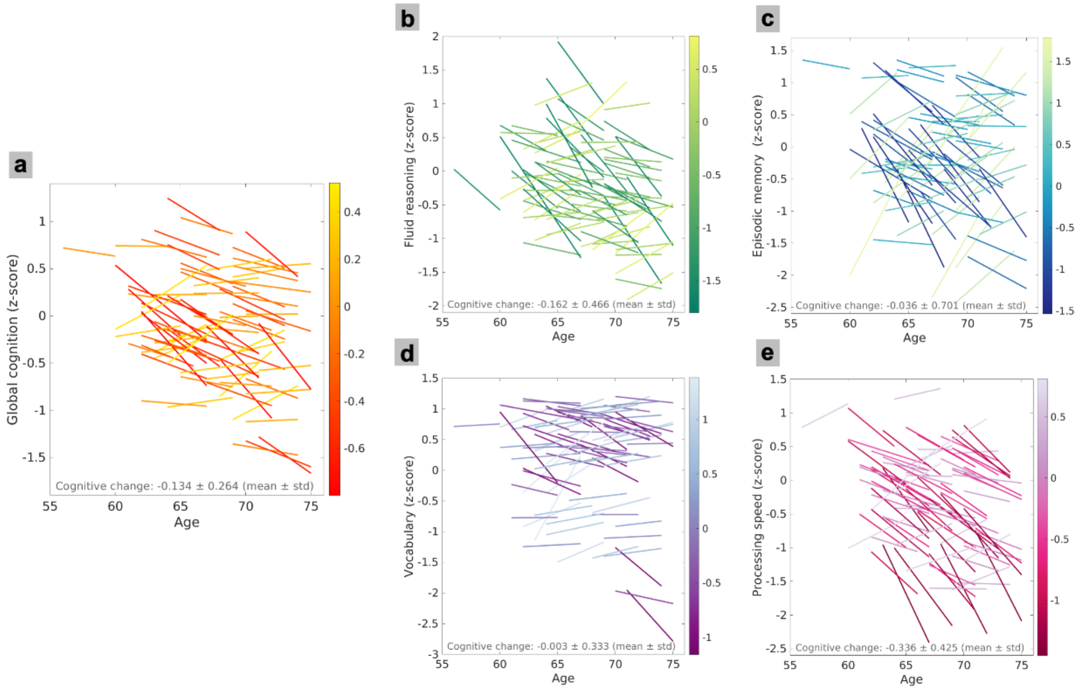


In the CogRes/RANN study, global cognition was calculated as the average of cognition z-scores in four cognitive domains (fluid reasoning, episodic memory, vocabulary, and processing speed). Participants showed cognitive decline in all four domains (fluid reasoning (**b**), episodic memory (**c**), vocabulary (**d**), and processing speed (**e**)) and global cognition (**a**). **a** Global cognition showed a longitudinal decline with a mean of -0.134 and standard deviation (SD) of 0.264. **b** Fluid reasoning showed a longitudinal decline with a mean of -0.162 and SD of 0.466. **c** Episodic memory showed a longitudinal decline with a mean of -0.036 and SD of 0.701. **d** Vocabulary showed a longitudinal decline with a mean of -0.003 and SD of 0.333. **e** Processing speed showed a longitudinal decline with a mean of -0.336 and SD of 0.425.

**Fig. S2.** Visualization of the centrality-scaled network-based amyloid-β pathology (NAP) scores and its regional contribution towards amyloid-cognition relationship.

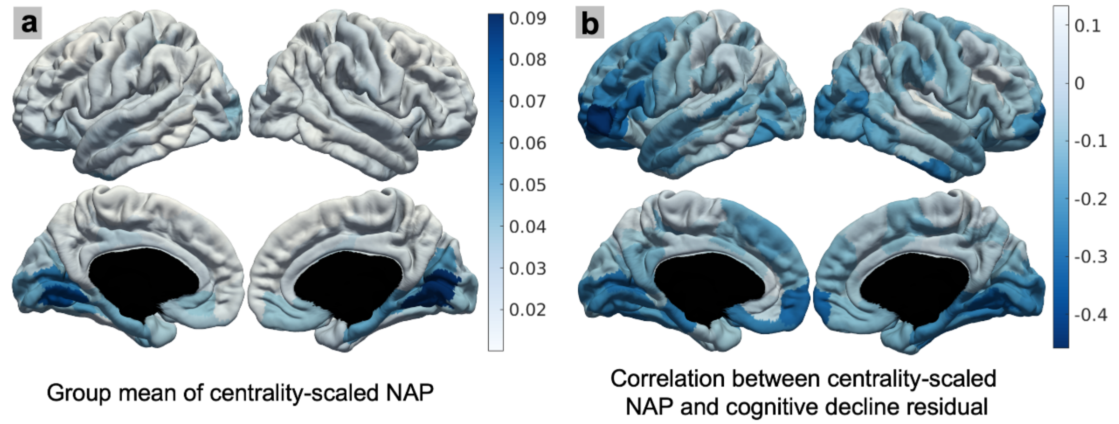


**a** Group averaged structural connectome (SC) centrality-scaled NAP scores. **b** Correlation coefficient between cognitive decline and SC centrality-scaled NAP scores across cortical regions of interest (Spearman partial correlation, controlling for age, sex, education, baseline cognition, and hemispherical mean cortical thickness).

**Fig. S3.** Performance comparison of functional connectivity (FC) based network-based amyloid-β pathology (NAP) measures with proportional thresholding versus regional amyloid-β pathology (RAP) measures.

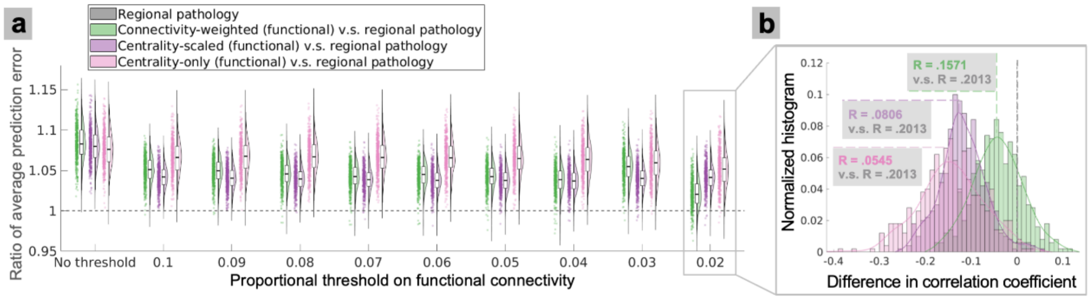


**a** *Ratio* (ratio of average prediction error) performance of FC-based NAP measures with different proportional thresholds. Keeping the top 2% of the connections achieved the lowest prediction error, with the best performance. **b** Correlation performance of FC-based NAP measures with the best performance model (proportional threshold of 2%). The best FC-based NAP model (connectivity-weighted NAP) had a predictive performance of median correlation coefficient R = 0.1571, which is still worse than the predictive performance of RAP measures (R = 0.2013).

**Fig. S4.** Longitudinal change in the ADNI Study cognition scores.

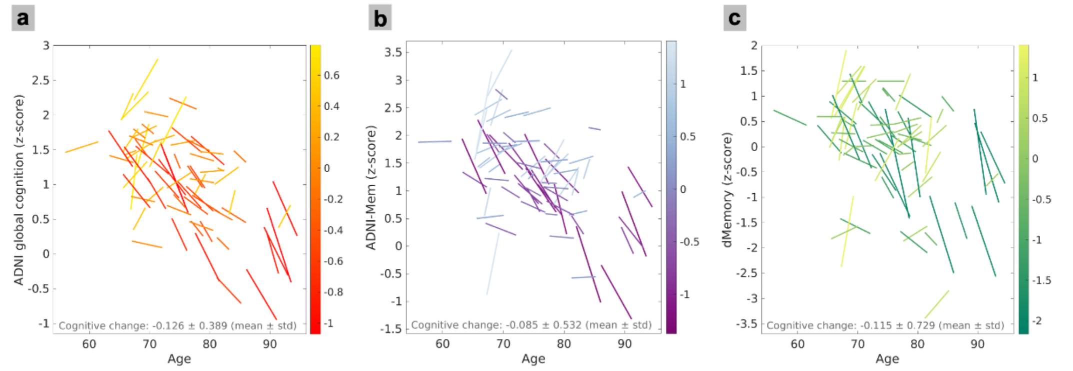

In the ADNI study, global cognition was calculated as the average cognition z-score in three cognitive domains (memory, executive functioning, and language). Participants showed cognitive decline in ADNI global cognition (ADNI-Gcog; **a**), ADNI memory score (ADNI-Mem; **b**), and delayed memory recall score (dMemory; **c**). **a** ADNI-Gcog showed a longitudinal decline with a mean of -0.126 and standard deviation (SD) of 0.389. **b** ADNI-Mem showed a longitudinal decline with a mean of -0.085 and SD of 0.532. **c** dMemory showed a longitudinal decline with a mean of -0.115 and SD of 0.729.

**Fig. S5.** Network-level correlation results in the relationship between pathology features in each brain network and longitudinal cognitive decline.

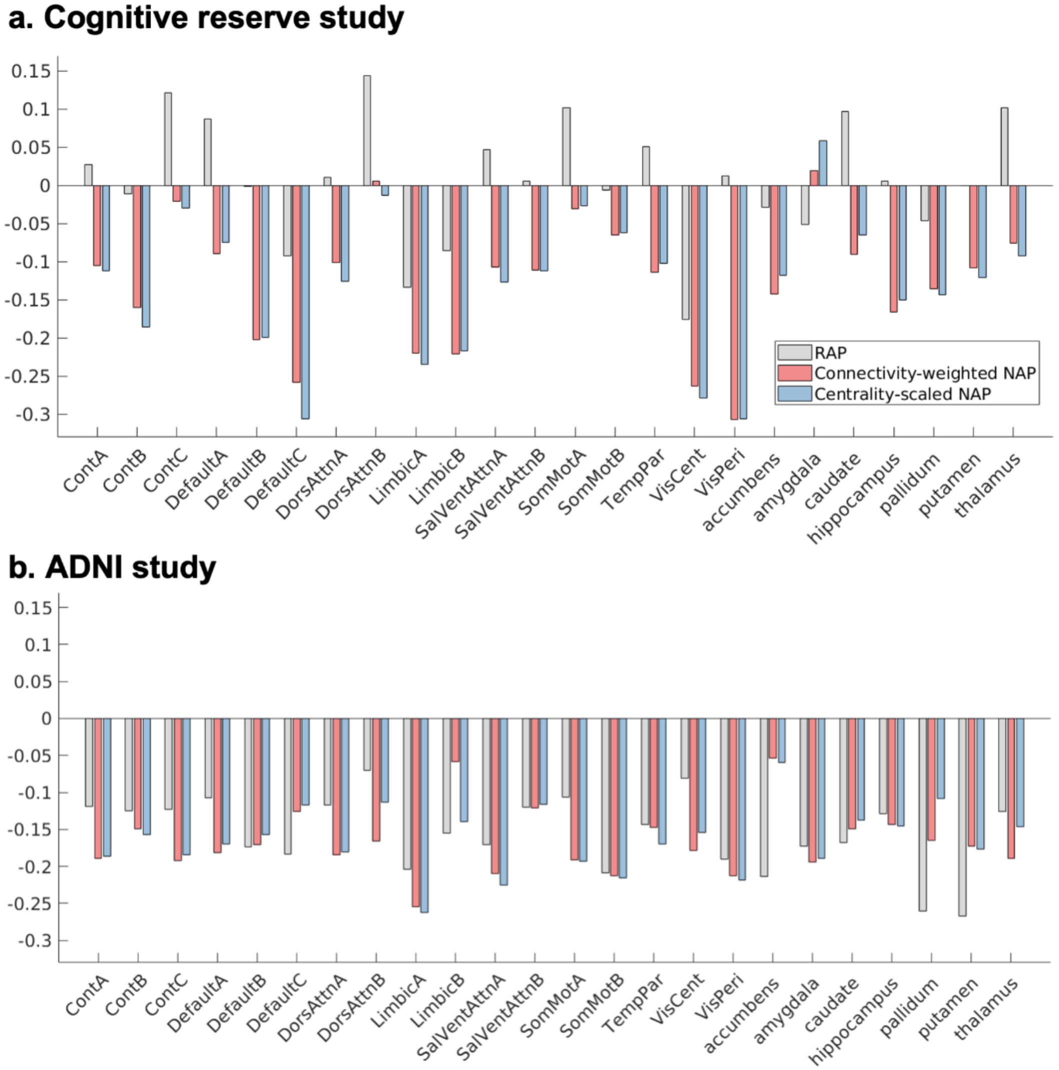


**a** For participants in the cognitive reserve and reference ability neural network (CogRes/RANN) longitudinal study, we performed Spearman partial correlation analysis between pathology features in each brain network and longitudinal cognitive changes, controlling for age, sex, education, baseline cognition, and mean cortical thickness of each hemisphere. Brain networks were defined by the Schaefer atlas. And pathology features were characterized by either regional amyloid-β pathology (RAP) or network-based amyloid-β pathology (NAP). **b** Same approach was performed for participants in the ADNI study. Compared to RAP measures, NAP scores demonstrated a stronger negative relationship with cognition change in both studies. Cognition was assessed using global cognition in CogRes/RANN study and dMemory in ADNI study.

**Table S1.** The neuropsychological assessments battery in the cognitive reserve and reference ability neural network (CogRes/RANN) longitudinal study and tests used in the present study for assessing cognition in domains of episodic memory, vocabulary, processing speed, and fluid reasoning. Global cognition was computed as an average of these four scores.

| Neuropsychological assessments | Episodic memory | Vocabulary | Processing speed | Fluid reasoning |
|---|---|---|---|---|
| Wechsler Adult Intelligence Scale (WAIS-III; Wechsler, 1997) | | | | |
| Letter-Number Sequencing | | | | |
| American National Adult Reading Test (AMNART; Wechsler, 1997) | | x | | |
| Selective Reminding Task (SRT) immediate recall (Buschke and Fuld, 1974) | x | | | |
| WAIS-III Matrix Reasoning (Wechsler, 1997) | | | | x |
| SRT delayed recall and delayed recognition (Buschke and Fuld, 1974) | x | | | |
| WAIS-III Digit Symbol (Wechsler, 1997) | | | x | |
| Trail-Making Test versions A and B (TMT-A/B; Reitan, 1978) | | | TMT-A | TMT-B |
| Controlled Word Association (C-F-L) and Category Fluency (animals; Benton et al., 1983) | | | | |
| Stroop Color Word Test (Golden, 1975) | | | x | |
| Wechsler Test of Adult Reading (WTAR; Holdnack, 2001) | | x | | |
| WAIS-III Vocabulary (Wechsler, 1997) | | x | | |
| WAIS-III Block Design (Wechsler, 1997) | | | | x |

**Table S2.** Regions in the amyloid-β neuropathological signatures of regional amyloid-β pathology (RAP), connectivity-weighted network-based amyloid-β pathology (NAP), or centrality-scaled NAP. The name, labels, and centroid coordinates of the regions were reported based on Schaefer atlas (200 Parcels and 17Networks version). Please be noted that the reported coordinates are in the volumetric MNI152 space (FSL-MNI152; 2mm version), however, analyses in this study are based on the surface parcellation and registration as its suggested to be optimal [Wu et al., 2018 Hum Brain Mapp].

| Regions | Labels | RAP signature | Connectivity-weighted NAP signature | Centrality-scaled NAP signature | Centroid coordinates in MNI152 space (RAS; right-anterior-superior) | | |
|---|---|---|---|---|---|---|---|
| | | | | | R | A | S |
| LH_VisCent_ExStr_1 | 1001 | | x | x | -26 | -78 | -14 |

| | | | | | | | |
|---|---|---|---|---|---|---|---|
| LH_VisCent_Striate_1 | 1003 | | x | | -6 | -92 | -4 |
| LH_VisPeri_ExStrInf_2 | 1008 | | x | x | -10 | -68 | -4 |
| LH_VisPeri_StriCal_1 | 1010 | | x | x | -12 | -70 | 8 |
| LH_DorsAttnA_TempOcc_1 | 1029 | x | | | -44 | -48 | -20 |
| LH_DorsAttnA_TempOcc_2 | 1030 | | x | x | -46 | -70 | -8 |
| LH_SalVentAttnA_FrOper_2 | 1043 | | x | x | -52 | 8 | 10 |
| LH_SalVentAttnB_Ins_1 | 1049 | | x | x | -34 | 20 | 6 |
| LH_LimbicB_OFC_2 | 1052 | x | | x | -10 | 36 | -20 |
| LH_ContA_PFClv_1 | 1062 | | | x | -42 | 40 | 16 |
| LH_ContA_PFCl_3 | 1065 | | | x | -42 | 6 | 44 |
| LH_ContB_PFCl_1 | 1069 | | x | | -40 | 18 | 50 |
| LH_ContB_PFClv_1 | 1070 | x | x | x | -42 | 50 | -6 |
| LH_DefaultA_PFCm_2 | 1081 | | x | x | -12 | 64 | -6 |
| LH_DefaultB_PFCv_2 | 1093 | | x | x | -32 | 42 | -14 |
| LH_DefaultB_PFCv_3 | 1094 | | x | x | -46 | 30 | -8 |
| LH_DefaultB_PFCv_4 | 1095 | | x | x | -52 | 22 | 8 |
| LH_DefaultC_PHC_1 | 1098 | | x | x | -26 | -32 | -18 |
| RH_VisCent_ExStr_1 | 2001 | | x | x | 28 | -68 | -12 |
| RH_VisCent_Striate_1 | 2003 | | x | | 12 | -92 | -6 |
| RH_VisCent_ExStr_3 | 2004 | | x | x | 30 | -94 | -4 |
| RH_VisPeri_ExStrInf_1 | 2007 | | x | x | 12 | -64 | -4 |
| RH_VisPeri_ExStrInf_2 | 2008 | | x | x | 16 | -46 | -2 |
| RH_DorsAttnA_ParOcc_1 | 2032 | | | x | 52 | -60 | 10 |
| RH_LimbicB_OFC_4 | 2060 | | x | x | 14 | 64 | -8 |
| RH_LimbicA_TempPole_1 | 2061 | | x | | 30 | 8 | -38 |
| RH_LimbicA_TempPole_2 | 2062 | | x | x | 46 | -12 | -34 |
| RH_ContB_PFClv_1 | 2077 | | x | x | 36 | 46 | -14 |
| RH_DefaultC_PHC_1 | 2096 | | | x | 28 | -36 | -14 |

**Table S3.** Comparison of the cognitive decline predictive performance of neuropathological signatures on CogRes/RANN dataset (* p < 0.05, ** p < 0.01).

| Cognitive decline measures | Regional amyloid-β pathology | Connectivity-weighted network amyloid-β pathology | Centrality-scaled network amyloid-β pathology |
|---|---|---|---|

| | | | |
|---|---|---|---|
| Global cognition | *r = 0.3290; p = 0.0035 (**)* | *r = 0.3713; p = 0.0009 (**)* | *r = 0.3716; p = 0.0009 (**)* |
| Fluid reasoning | *r = 0.2037; p = 0.0756* | *r = 0.1542; p = 0.1805* | *r = 0.1932; p = 0.0924* |
| Episodic memory | *r = 0.2937; p = 0.0095 (**)* | *r = 0.2855; p = 0.0118 (*)* | *r = 0.2768; p = 0.0148 (*)* |
| Vocabulary | *r = 0.0621; p = 0.5917* | *r = 0.1796; p = 0.1181* | *r = 0.1726; p = 0.1333* |
| Processing speed | *r = 0.1600; p = 0.1644* | *r = 0.1952; p = 0.0890* | *r = 0.1989; p = 0.0829* |

1. Baker, J. E. *et al.* Cognitive impairment and decline in cognitively normal older adults with high amyloid-β: A meta-analysis. *Alzheimer's & Dementia: Diagnosis, Assessment & Disease Monitoring* 6, 108–121 (2017).
